# VYD2311 is a promising candidate for passive immunization against COVID-19 in immunocompromised individuals

**DOI:** 10.64898/2026.03.31.715419

**Authors:** Ian A. Mellis, Madeline Wu, Kristin Daniel, Hsiang Hong, Yicheng Guo, David D. Ho

**Affiliations:** Aaron Diamond AIDS Research Center, Columbia University Vagelos College of Physicians and Surgeons, New York, NY, USA; Department of Pathology and Cell Biology, Columbia University Vagelos College of Physicians and Surgeons, New York, NY 10032, USA; Department of Microbiology, National Taiwan University College of Medicine, Taipei, Taiwan; Division of Infectious Diseases, Department of Medicine, Columbia University Vagelos College of Physicians and Surgeons, New York, NY, USA; Pandemic Research Alliance unit at the Wu Center for Pandemic Research, Columbia University Vagelos College of Physicians and Surgeons, New York, NY 10032, USA

**Author notes:** Correspondence (I.A.M.). equal contribution (Ian A. Mellis, Madeline Wu, Kristin Daniel, Yicheng Guo).

## Abstract

For millions of immunocompromised individuals, vaccines may not elicit adequate protection from infections, so alternative strategies for pre-exposure prophylaxis are essential. There is only one non-vaccine product authorized in the U.S. as pre-exposure prophylaxis against COVID-19: the monoclonal antibody pemivibart. We previously showed that pemivibart had lower neutralizing activity *in vitro* against many recent dominant SARS-CoV-2 variants, such as KP.3.1.1, NB.1.8.1, and LP.8.1.1, than it had against JN.1, which was dominant when the antibody was first authorized. The manufacturer of pemivibart (Invivyd) recently initated clinical testing of a new monoclonal antibody derived from pemivibart, VYD2311, but there are no available studies of the activity of VYD2311 against dominant and emerging SARS-CoV-2 variants. Here, using pseudovirus neutralization assays, we measured the neutralizing activity of laboratory-synthesized VYD2311 and pemivibart against dominant and emerging SARS-CoV-2 variants, including XFG, NB.1.8.1, and the genetically distant BA.3.2.2. We found that VYD2311 potently neutralized all tested variants *in vitro*, dramatically more so than pemivibart. Combined with interpretation of earlier clinical trials of a parental antibody product, we conclude that VYD2311 is a promising candidate for passive immunoprophylaxis against COVID-19, particularly for those who do not respond well to vaccination.

## Main Text

For millions of immunocompromised individuals, vaccines may not elicit adequate protection from infections, so alternative strategies for pre-exposure prophylaxis are essential. There is only one non-vaccine product authorized in the U.S. as pre-exposure prophylaxis against coronavirus disease 2019 (COVID-19): pemivibart. Pemivibart is a monoclonal antibody targeting the severe acute respiratory syndrome coronavirus 2 (SARS-CoV-2) spike protein receptor-binding domain class 1/4 region, which was authorized in March 2024, when the JN.1 variant was dominant^1^. We showed that pemivibart had lower neutralizing activity *in vitro* against recent SARS-CoV-2 variants, such as KP.3.1.1, NB.1.8.1, and LP.8.1.1, than against JN.1^2,3^. In 2024, Invivyd initiated clinical testing of a new monoclonal antibody, VYD2311, and subsequently registered a phase 3 clinical trial to test it as pre-exposure prophylaxis against COVID-19^4,5^. VYD2311 was derived from pemivibart, containing six mutations (three in the heavy chain and three in the light chain; **Figure S1**). In a patent, Invivyd reported that VYD2311 had high neutralizing potency against recent JN.1 subvariants, including KP.3.1.1 and LP.8.1, but they did not report results for currently dominant and expanding variants, such as XFG, NB.1.8.1, and the genetically distant BA.3.2.2 (**Figure S2**)^6^. It is essential to monitor the efficacy of antibody products in the setting of evolving viral variants.

Using pseudovirus neutralization assays, we measured the neutralizing activities of versions of VYD2311 and pemivibart synthesized in our laboratory against recent, dominant, and expanding SARS-CoV-2 variants: D614G (representing the ancestral strain), JN.1, KP.3.1.1, LP.8.1.1, NB.1.8.1, XFG, and BA.3.2.2 (**Figure 1**)^3^. The 50% inhibitory concentrations (IC_50_) of laboratory-synthesized VYD2311 against JN.1, XFG, BA.3.2.2, and NB.1.8.1 were approximately 0.004, 0.002, 0.013, and 0.126 µg/mL, respectively. The 90% inhibitory concentrations (IC_90_) of VYD2311 against JN.1, XFG, BA.3.2.2, and NB.1.8.1 were approximately 0.029, 0.030, 0.126, and 1.05 µg/mL. Comparing laboratory-synthesized VYD2311 to laboratory-synthesized pemivibart, the IC_50_s against JN.1, XFG, BA.3.2.2, and NB.1.8.1 were approximately 27, 51.5, 31.54, and 123.6-fold lower, with even larger factor changes in IC_90_ (**Figure S3**).

**Figure 1:**
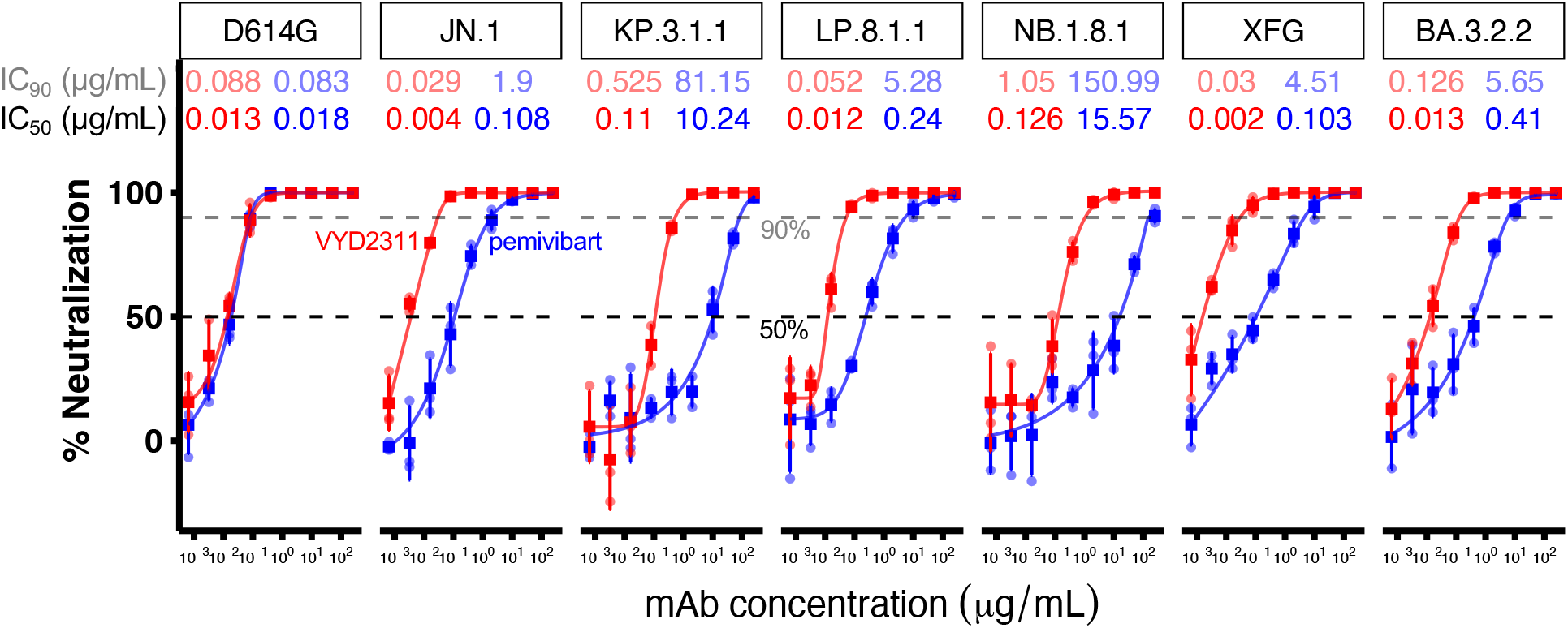
Neutralizing activity of research-grade VYD2311 and research-grade pemivibart. Percent neutralization of the indicated pseudotyped viruses by VYD2311 (red) and pemivibart (blue). Dots indicate technical replicates, squares indicate mean, I bars indicate standard deviation. 5-fold serial dilutions from a maximum tested concentration of 250 µg/mL, nine doses tested. Dashed lines indicate 50% and 90% neutralizations. The indicated inhibitory concentrations (ICs) of each antibody at 50% and 90% neutralization levels against each variant are annotated above each pair of curves.

Pemivibart is authorized to be administered intravenously every 3 months at a dose of 4500 mg and was reported to have a trough serum concentration of approximately 175 µg/mL. Adintrevimab, the parental molecule to pemivibart, was tested in Phase 1 trials at 300 mg or 600 mg intramuscular doses or at a 500 mg intravenous dose. The mean serum adintrevimab concentration 90 days post-administration in a peer-reviewed study was >10 µg/mL for all three administration methods^7^. The VYD2311 patent also reports Phase 1/2 testing of 1000 mg intramuscular, 1250 mg subcutaneous, and 2000 mg or 4500 mg intravenous VYD2311 doses, which led to mean serum concentrations of >10 µg/mL at 90 days post-administration (or 45 days for subcutaneous dosing)^6^. Therefore, if VYD2311 pharmacokinetics are confirmed and similar to adintrevimab, then at the reported doses VYD2311 could circulate in the blood at higher than the IC_99_ values of all tested variants for at least 90 days (**Figure S3**). In summary, VYD2311 is a promising investigational product for the prevention and treatment of COVID-19.

