## Supplementary Appendix for "VYD2311 is a promising candidate for passive immunization against COVID-19 in immunocompromised individuals"

### Supplementary Figures and Tables

**Figure S1: Amino acid sequences of studied antibodies**

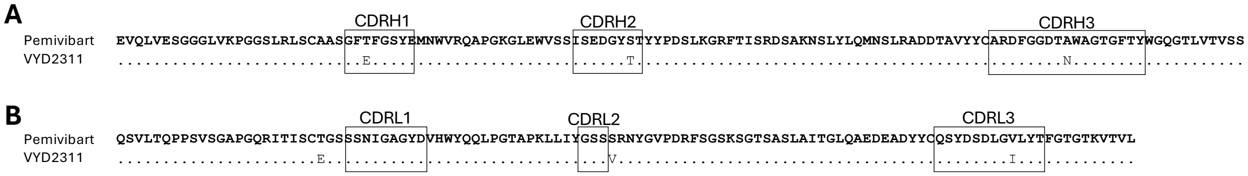

**Figure S1: Amino acid sequences of studied antibodies**. Multiple sequence alignment of the VH and VL sequence variable regions for VYD2311 and pemivibart. VYD2311 sequence extracted from patent WO2026010909A1 (Invivyd) and aligned here[1]. A: heavy chains. B: light chains. The CDRs in this alignment were defined according to the IMGT scheme.

**Figure S2: SARS-CoV-2 variant spike protein mutations**

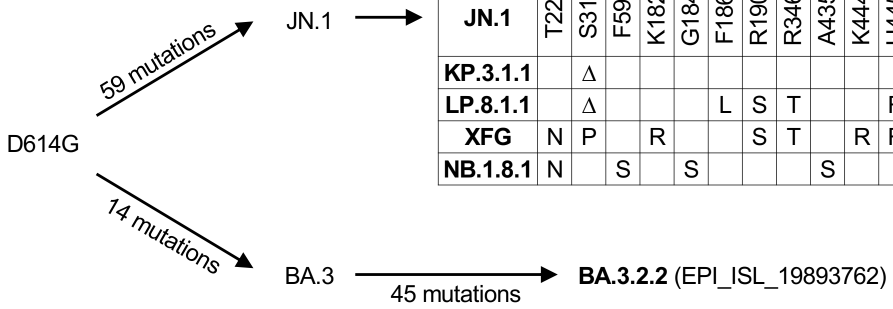

| | NTD | | | | | | | RBD | | | | | | | | SD1 | S2 | | |
|---|---|---|---|---|---|---|---|---|---|---|---|---|---|---|---|---|---|---|---|
| **JN.1** | T22 | S31 | F59 | K182 | G184 | F186 | R190 | R346 | A435 | K444 | H445 | F456 | K478 | N487 | Q493 | T572 | K679 | K1086 | V1104 |
| **KP.3.1.1** | | Δ | | | | | | | | | | L | | | E | | | | L |
| **LP.8.1.1** | | Δ | | | | L | S | T | | | R | L | | | E | | R | R | L |
| **XFG** | N | P | | R | | | S | T | | R | R | L | | D | E | I | | | |
| **NB.1.8.1** | N | | S | | S | | | | S | | | L | I | | E | | | | |

**Figure S2: SARS-CoV-2 variant spike protein mutations**. Selected phylogenetic and genetic relationships between SARS-CoV-2 variants of interest. The ancestral strain is represented by D614G. JN.1 is the parental variant of many dominant and recent subvariants. Spike mutations relative to JN.1 are shown for the indicated subvariants. BA.3.2.2 is an emerging variant descended from BA.3. The GISAID sequence identifier corresponding to the BA.3.2.2 spike protein sequence is indicated. Δ: deletion.

**Figure S3: Comparison of the neutralizing activities of research-grade VYD2311 and research-grade pemivibart**

| | IC$_{50}$ (µg/mL) | | factor change | IC$_{80}$ (µg/mL) | | factor change | IC$_{90}$ (µg/mL) | | factor change | IC$_{99}$ (µg/mL) | | factor change |
|---|---|---|---|---|---|---|---|---|---|---|---|---|
| | pemivibart | VYD2311 | | pemivibart | VYD2311 | | pemivibart | VYD2311 | | pemivibart | VYD2311 | |
| D614G | 0.02 | 0.01 | 1.38 | 0.06 | 0.05 | 1.15 | 0.08 | 0.09 | 0.94 | 0.21 | 0.46 | 0.47 |
| JN.1 | 0.11 | 0.004 | 27 | 0.69 | 0.02 | 42.81 | 1.9 | 0.03 | 65.52 | 37.47 | 0.1 | 390.28 |
| KP.3.1.1 | 10.25 | 0.11 | 93.14 | 45.47 | 0.29 | 156.26 | 81.15 | 0.53 | 154.58 | >250 | 2.73 | >91.74 |
| LP.8.1.1 | 0.24 | 0.01 | 20 | 1.65 | 0.03 | 54.9 | 5.28 | 0.05 | 101.52 | 164.2 | 0.32 | 517.99 |
| NB.1.8.1 | 15.57 | 0.13 | 123.6 | 91.02 | 0.47 | 193.66 | 150.99 | 1.05 | 143.52 | >250 | 8.71 | >28.69 |
| XFG | 0.1 | 0.002 | 51.5 | 1.59 | 0.01 | 144.64 | 4.51 | 0.03 | 150.33 | 37.04 | 0.42 | 88.4 |
| BA.3.2.2 | 0.41 | 0.01 | 31.54 | 2.64 | 0.06 | 41.87 | 5.65 | 0.13 | 44.85 | 71.43 | 0.71 | 100.04 |

**Figure S3: Comparison of the neutralizing activities of research-grade VYD2311 and research-grade pemivibart**. 50%, 80%, 90%, and 99% inhibitory concentrations of each antibody (IC$_{50}$, IC$_{80}$, IC$_{90}$, IC$_{99}$, µg/mL) against the indicated variants are shown. 5-fold serial dilutions from a maximum tested concentration of 250 µg/mL, nine doses tested. Darker shades of orange indicate lower IC. Darker shades of green indicate larger fold-differences in the indicated inhibitory concentrations between VYD2311 and pemivibart.

### Supplementary Methods

#### Cell lines

Vero-E6 (CRL-1586) cells and HEK293T (CRL-3216) cells were obtained from ATCC and cultured at 37°C in an atmosphere of 5% $CO_2$ in Dulbecco's Modified Eagle Medium (DMEM) + 10% fetal bovine serum (FBS) + 1% penicillin-streptomycin. Expi293F (A14527) cells were purchased from ThermoFisher and maintained in Expi293 expression medium per the manufacturer's instructions. Vero-E6 cells are derived from African green monkey kidneys. HEK293T cells and Expi293 cells are human female in origin.

#### Plasmid generation

Research-grade pemivibart heavy chain and light chain expression plasmids were previously constructed by our laboratory as previously described[2]. To generate VYD2311 antibody expression plasmids, mutations were made to our pemivibart heavy chain and light chain constructs using the QuikChange II XL and QuikChange Multi site-directed mutagenesis kits (Agilent). All plasmids were verified by whole plasmid sequencing prior to further use.

#### Protein expression and purification

For each antibody of interest, the corresponding heavy chain and light chain plasmids were transfected into Expi293F cells at a 1:1 ratio, and then the supernatants were collected after five days. The antibodies were purified with Protein A Sepharose (Cytiva) following the manufacturer's instructions. Molecular weight and purity were confirmed by SDS-PAGE protein electrophoresis to use.

#### Pseudovirus production

VSV-based SARS-CoV-2 pseudoviruses, in which the native VSV glycoprotein was replaced by SARS-CoV-2 variant spikes, were produced as previously described. Briefly, plasmids containing the appropriate spike were transfected into HEK293T cells with PEI-MAX. After 24 hours, VSV-G pseudotyped ΔG-luciferase (G*ΔG-luciferase, Kerafast) was added and then washed with DPBS three times before being cultured in fresh medium for another 24 hours. Anti-VSVG (anti-I1) antibody was added to deplete non-pseudotyped viruses. Pseudoviruses were then harvested, centrifuged, and then aliquoted and stored at -80°C.

#### Pseudovirus neutralization assays

Each SARS-CoV-2 pseudovirus was titrated to standardize infectious dose before use in neutralization assays. 5-fold serial dilutions of monoclonal antibodies (mAbs) were added in 96-well plates, starting at the concentrations indicated in each figure legend. Next, pseudoviruses were added and incubated at 37 °C for 1 hour. In each plate, wells containing only pseudoviruses were included as controls. 40,000 Vero-E6 cells were then added per well and incubated at 37 °C for 16 hours. Promega Luciferase Assay System (E4550) was used for lysis and luciferase activity measurements on a Tecan Infinite® 200 PRO using i-control™ software v.3.9.1.0, following the manufacturer's instructions.

### Quantification and statistical analysis

The mAb concentration that 50% neutralizes ($IC_{50}$) and 90% neutralizes ($IC_{90}$) each viral variant was calculated using five-parameter log-logistic dose-response curve fitting with the drda package v2.0.5 in R v4.3.2.

### Author Contributions

The study was conceptualized by I.A.M., Y.G., and D.D.H. Experiments were conducted and data analyzed by I.A.M., M.W., H.H., and K.D. Project management was handled by I.A.M. The results were interpreted and the manuscript was written by I.A.M., M.W., K.D., Y.G., and D.D.H. All contributing authors have reviewed and endorsed the manuscript. I.A.M, M.W., K.D., and Y.G. contributed equally.

### Declaration of Interests

D.D.H. co-founded TaiMed Biologics and RenBio, and he serves as a consultant for WuXi Biologics and Brii Biosciences and is a board director at Vicarious Surgical. The remaining authors declare no conflicts of interest. None of the authors hold equity in, receive payments from, or consult for Invivyd.

### Acknowledgements

This study was supported by funding from the NIH SARS-CoV-2 Assessment of Viral Evolution (SAVE) Program (subcontract no. 0258-A700-4609 under federal contract no. 75N93021C00014 and the Gates Foundation (project INV019355) to D.D.H. and internal startup funding UR014016 from Columbia University to Y.G. We thank all who contributed their data to the Global Initiative on Sharing All Influenza Data (GISAID).
